## Supplemental figures for "Dysfunctional S1P/S1PR1 signaling in the dentate gyrus drives vulnerability of chronic pain-related memory impairment"

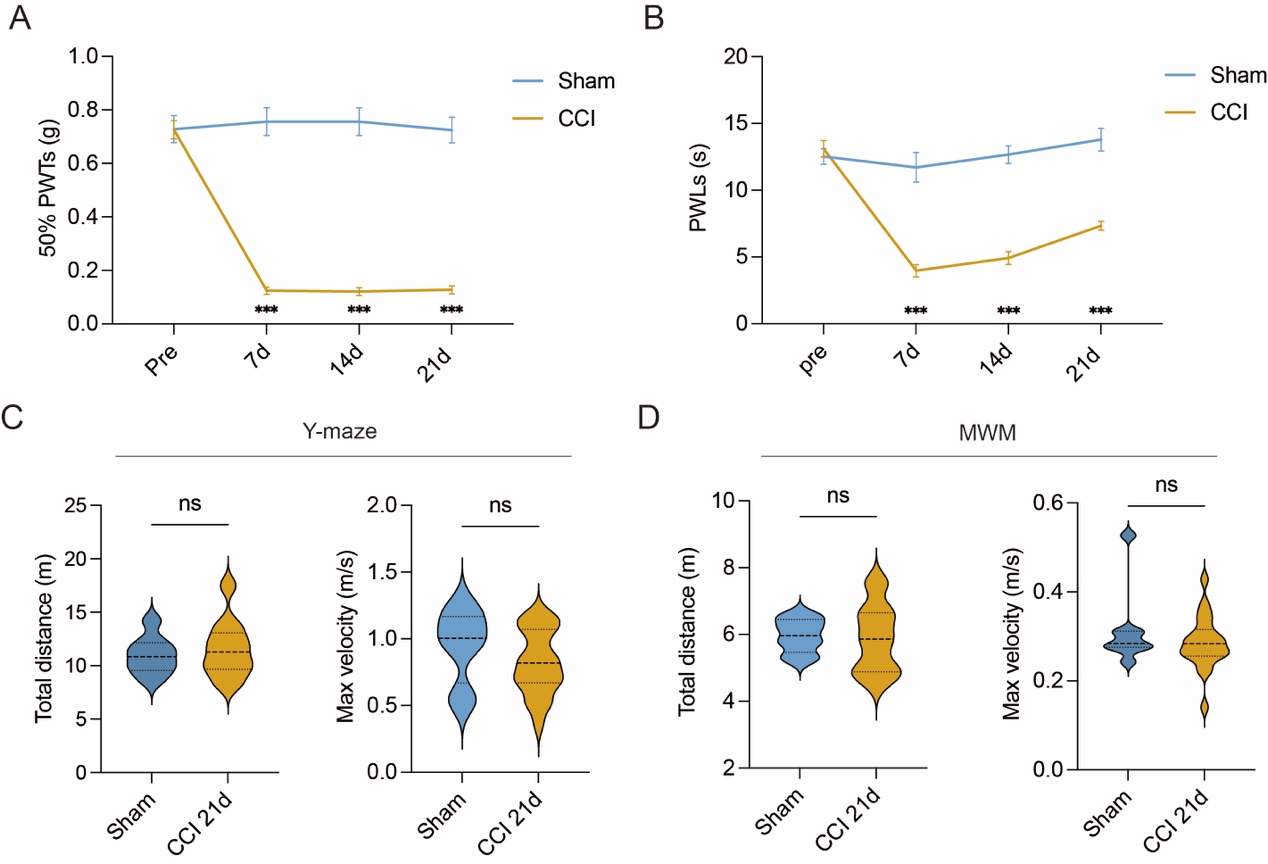

**Supplemental Figure 1. Behavioral assays of nociception and locomotor activity.** **(A-B)** 50% PWTs and PWLs of Sham- and CCI-treated mice of baseline and at 7d, 14d, 21d post CCI. **(C)** Quantitative summary of Y-maze test showing total distance traveled and max velocity in the novel arm in Sham- and CCI-Chronic mice (22d post CCI, n = 8-21). **(D)** Quantitative summary of MWM test showing total distance traveled and max velocity in the target quadrant in Sham- and CCI-Chronic mice (28d post CCI, n = 8-21). Data were analyzed by two-way ANOVA with post hoc Tukey’ s multiple comparisons test between groups **(A and B)** or two-tailed unpaired Student’s t-test **(C and D)**. All data are presented as the mean ± s.e.m. ns, not significant; ****p* < 0.001. CCI, chronic constrictive injury; PWTs, paw withdrawal thresholds; PWLs, paw withdrawal latencies; d, day.

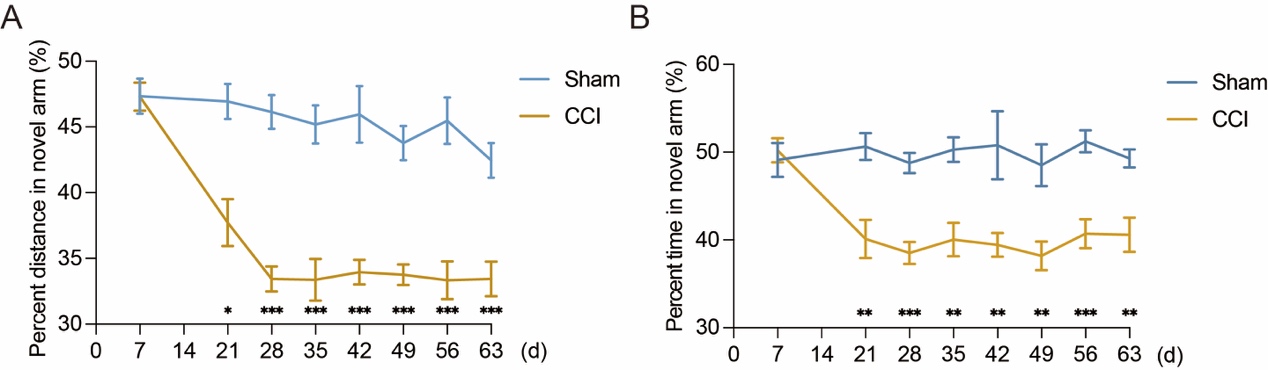

**Supplemental Figure 2. Chronic pain induced memory impairment lasts at least to 63d after CCI (A-B)** Quantitative summary of Y-maze test showing percent distance traveled and percent time spent in the novel arm in Sham- and CCI-treated mice at 7d, 21d, 28d, 35d, 42d, 49d, 56d, 63d after CCI (n = 10-20). Data were analyzed by two-way ANOVA with post hoc Tukey’ s multiple comparisons test between groups. All data are presented as the mean ± s.e.m. **p* < 0.05; ***p* < 0.01; ****p* < 0.001. CCI, chronic constrictive injury; d, day.

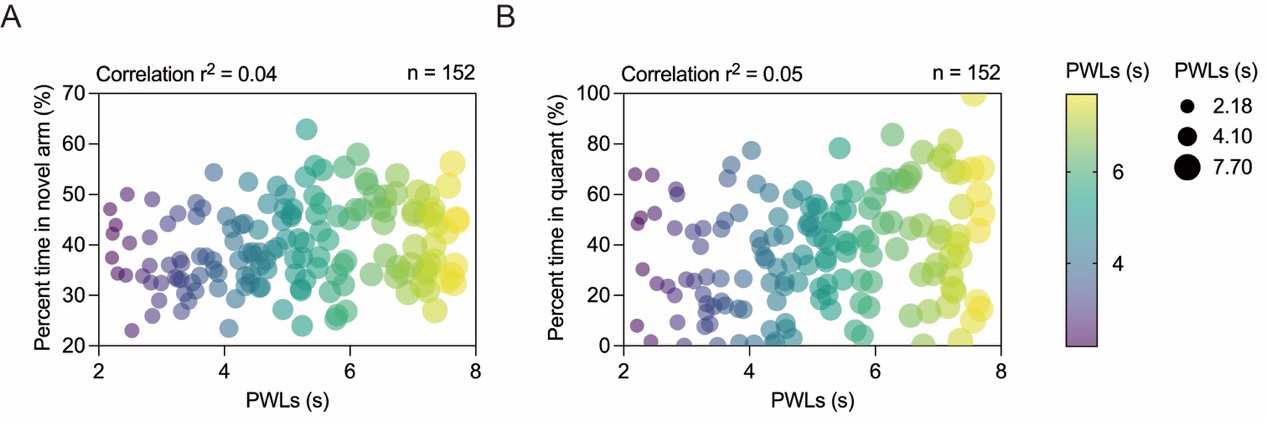

**Supplemental Figure 3. Susceptibility or insusceptibility to chronic pain induced memory impairment is irrelevant to pain threshold of CCI-treated mice (A-B)** Correlation analysis of PWLs with percent time in novel arm (r^2^=0.04, n = 152) and percent time in quadrant (r^2^=0.05, n = 152). Linear regression followed by a goodness-of-fit measure of R-squared (r^2^) was used to determine the correlation. PWLs, paw withdrawal latencies.

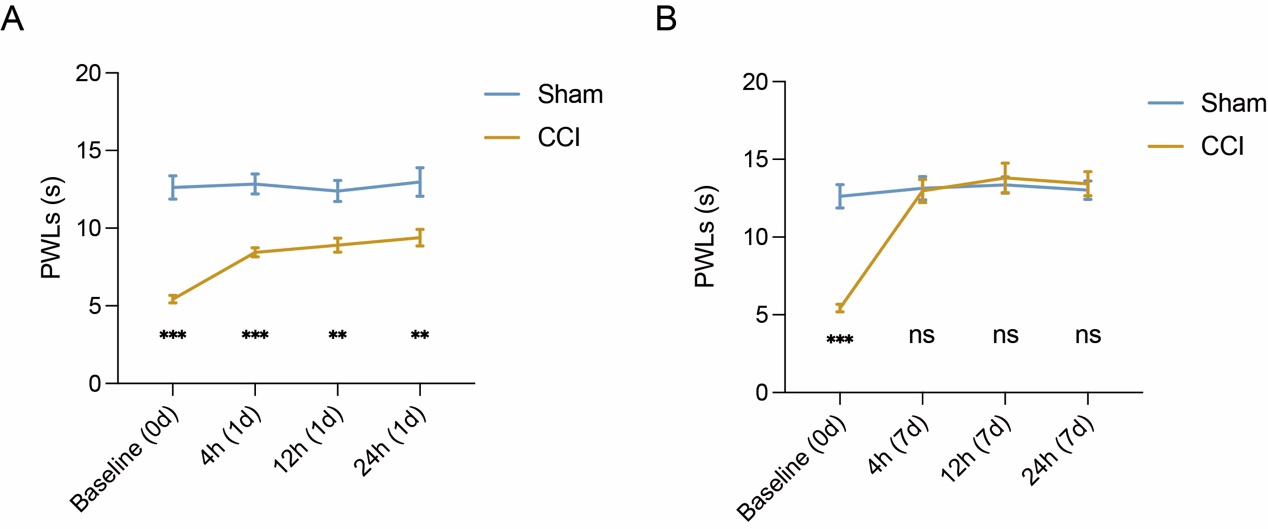

**Supplemental Figure 4. Analgesic effects of meloxicam on neuropathic pain of CCI. (A)** The analgesic effects of meloxicam on neuropathic pain of CCI 4h, 12h and 24h post a single dose of meloxicam (10mg/kg, *i.p.*). **(B)** The analgesic effects of meloxicam on neuropathic pain of CCI 4h, 12h and 24h on day 7 post intraperitoneal injection of meloxicam once daily (10mg/kg, *i.p.*). Data were analyzed by two-way ANOVA with post hoc Tukey’ s multiple comparisons test between groups. All data are presented as the mean ± s.e.m. ***p* < 0.01; ****p* < 0.001; ns, not significant. CCI, chronic constrictive injury; PWLs, paw withdrawal latencies; d, day; h, hour.

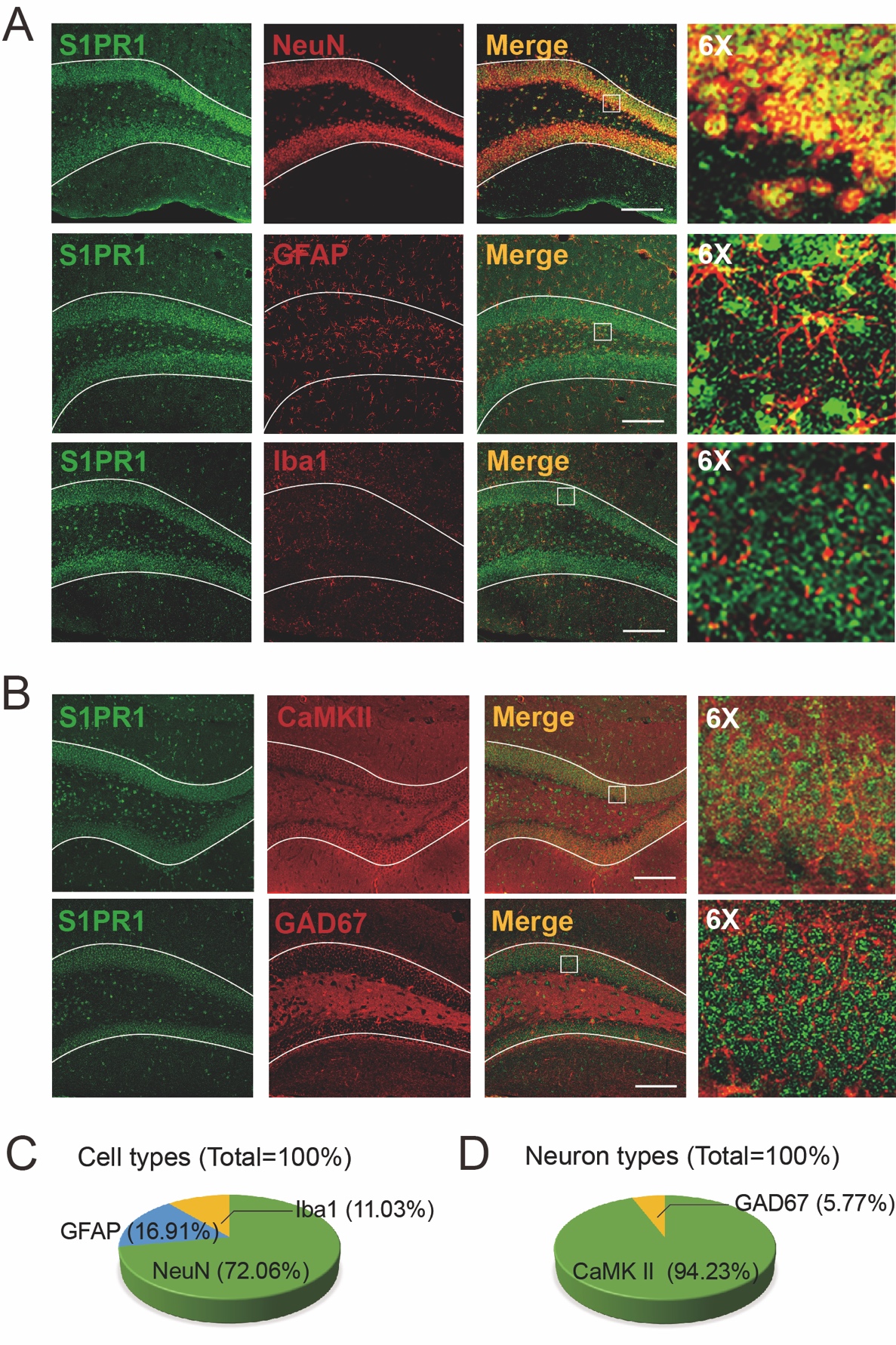

**Supplemental Figure 5. Characterization of expression profile of S1PR1 in the DG. (A-B)** Characterization of S1PR1 expression in different cell types **(A)** and neuron types **(B)** (Scale bar, 100 μm). **(C)** S1PR1 was highly coexpressed with NeuN and sparsely coexpressed with GFAP or Iba1 (Left, n = 4). **(D)** S1PR1 expression was found mostly in CaMKII+ neurons and sparsely in GAD67+ neurons (Right, n = 4).

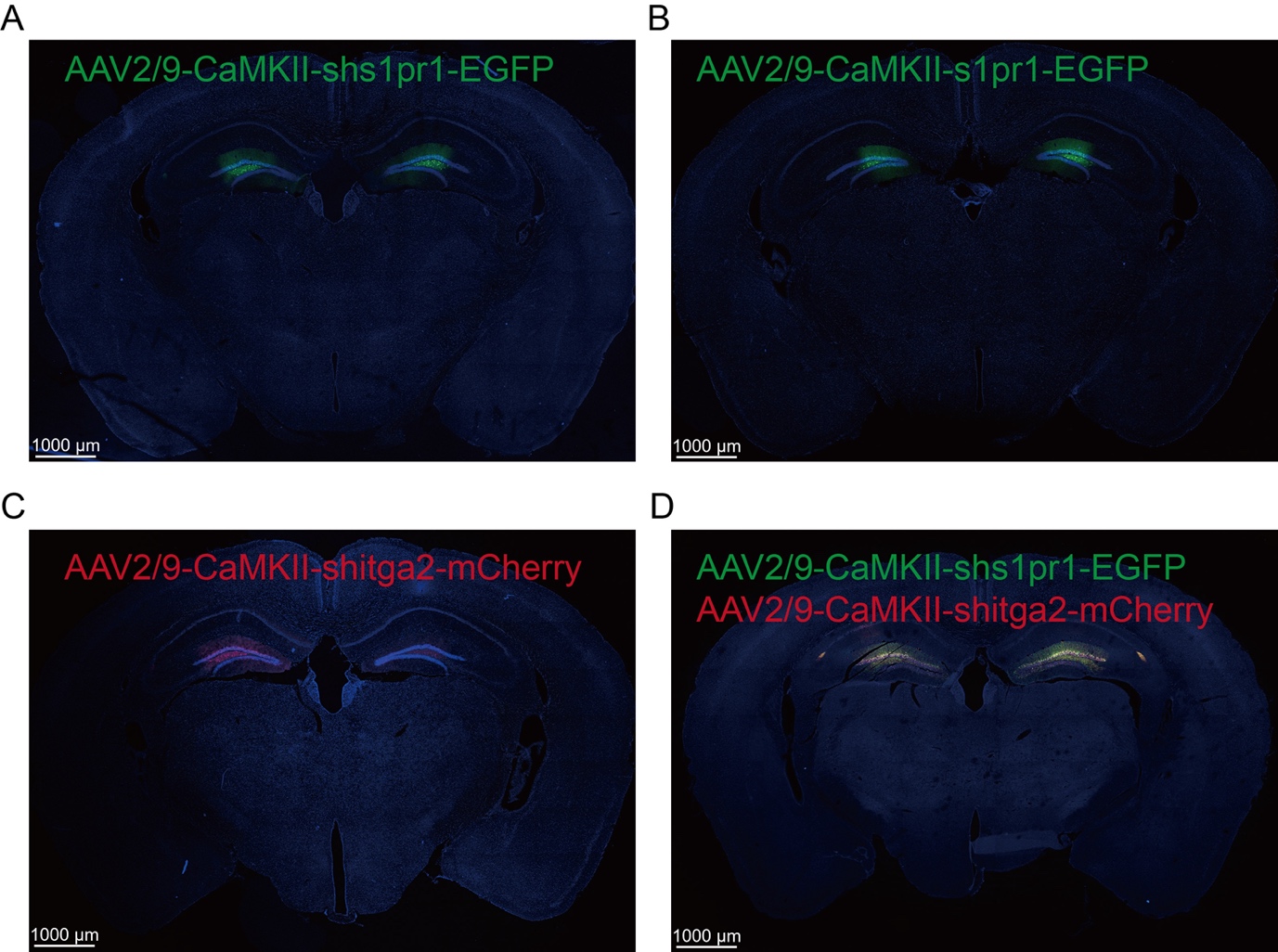

**Supplemental Figure 6. Zoomed-out images of the brain to show the precision of the virus injection (Scale bar, 1000 μm).**

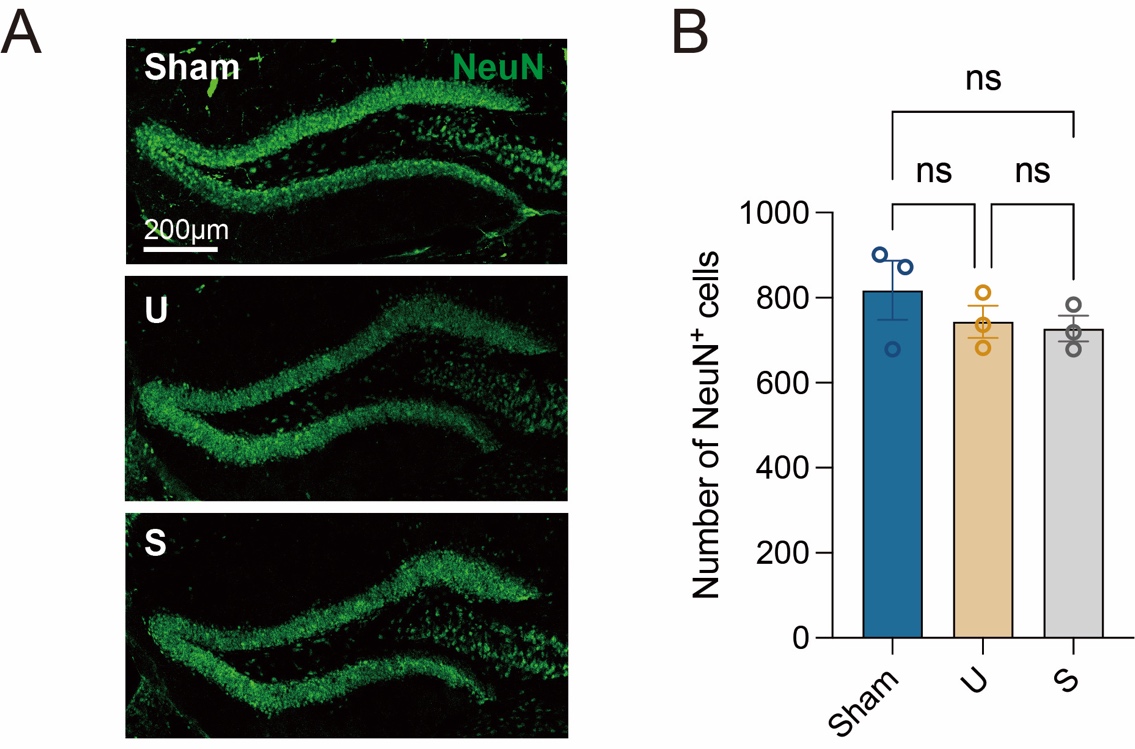

**Supplemental Figure 7. The number of neurons in the hippocampal dentate gyrus.** **(A)** Representative confocal images showing neurons in the DG in Sham, U and S mice (Scale bar, 200 μm). **(B)** Bar graph indicates the number of neurons in the DG (n = 3). Data were analyzed by one-way analysis of variance (one-way ANOVA), followed by post hoc Tukey’s multiple comparisons between multiple groups when appropriate. All data are presented as the mean ± s.e.m. ns, not significant. DG, dentate gyrus; U, unsusceptible; S, susceptible.

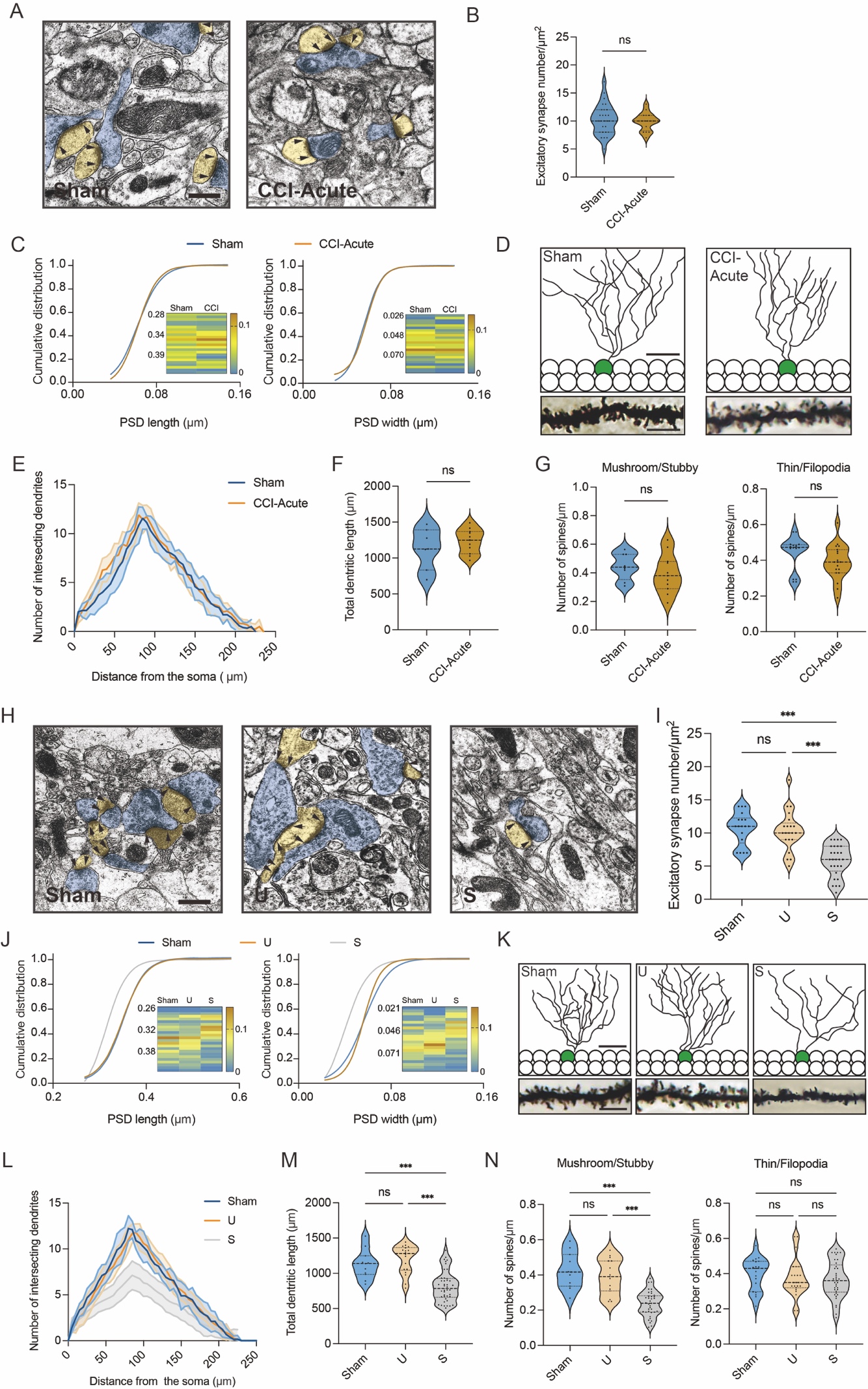

**Supplemental Figure 8. Susceptible mice exhibit altered excitatory synaptic plasticity in the hippocampal dentate gyrus.** **(A)** Representative TEM images of synapses in the DG in Sham and CCI-Acute mice (7d post CCI). Blue indicates presynaptic site and yellow indicates postsynaptic sites of excitatory synapses, respectively. Synaptic densities are bracketed by arrows (Scale bar, 500 nm). **(B)** Mean number of excitatory synapses per μm^2^ of DG in Sham and CCI-Acute mice (7d post CCI, n = 18-24 from 4 mice/group). **(C)** Cumulative distribution plots for the lengths and widths of postsynaptic density in the DG in Sham and CCI-Acute mice (n = 102-156 from 4 mice/group). **(D)** Representative Golgi-staining images of dendritic spine morphology from the DG in Sham and CCI-Acute mice (7d post CCI, Scale bar, top: 50 μm; bottom:10 μm). **(E)** The number of intersections of all dendritic branches in Sham and CCI-Acute mice (n = 15-18 from 4 mice/group). **(F)** Violin plots indicate the total dendritic length. **(G)** Violin plots indicate the number of mushroom/stubby type dendritic spines (left), and the number of thin/filopodia type dendritic spines (right) in Sham and CCI-Acute mice (7d post CCI, n = 15-18 from 4 mice/group). **(H)** Representative TEM images of synapses in the DG in Sham and CCI-Chronic mice (27d post CCI). Blue indicates presynaptic site and yellow indicates postsynaptic sites of excitatory synapses, respectively. Synaptic densities are bracketed by arrows (Scale bar, 500 nm). **(I)** Mean number of excitatory synapses per μm^2^ of DG in Sham, unsusceptible and susceptible mice (n = 18-24 from 4 mice/group). **(J)** Cumulative distribution plots for the lengths and widths of postsynaptic density in the DG in Sham, unsusceptible and susceptible mice (n = 102-156 from 4 mice/group). **(K)** Representative Golgi-staining images of dendritic spine morphology from the DG in Sham, unsusceptible and susceptible mice (Scale bar, top:50 μm; bottom:10μm). **(L)** The number of intersections of all dendritic branches in Sham, unsusceptible and susceptible mice (n = 15-18 from 4 mice/group). **(M)** Violin plots indicate the total dendritic length. **(N)** Violin plots indicate the number of mushroom/stubby type dendritic spines (left), and the number of thin/filopodia type dendritic spines (right) in Sham, unsusceptible and susceptible mice (n = 15-18 from 4 mice/group). Data were analyzed by unpaired t test or one-way analysis of variance (one-way ANOVA), followed by post hoc Tukey’s multiple comparisons between multiple groups when appropriate. All data are presented as the mean ± s.e.m. ns, not significant; ****p* < 0.001. CCI, chronic constrictive injury; d, day; DG, dentate gyrus; U, unsusceptible; S, susceptible; TEM, transmission electron microscope.

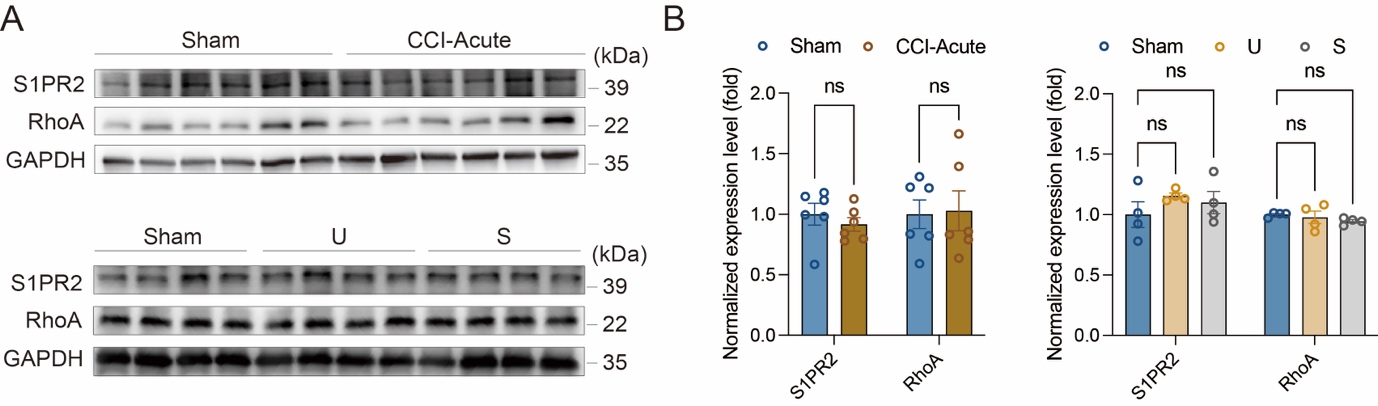

**Supplemental Figure 9. Expression levels of S1PR2 and RhoA in Sham and CCI animals. (A)** Example Western bands showing expression of S1PR2 and RhoA in DG lysates from Sham and CCI-Acute mice (7d post CCI), Sham and CCI-Chronic mice (Unsusceptible and Susceptible populations, 21d post CCI). **(B)** Densitometric comparison of the average expression of S1PR2 and RhoA (n = 6 for CCI-Acute mice, n = 4 for CCI-Chronic mice). Data were analyzed by unpaired t test or one-way analysis of variance (one-way ANOVA), followed by post hoc Tukey’s multiple comparisons between multiple groups when appropriate. All data are presented as the mean ± s.e.m. ns, not significant. CCI, chronic constrictive injury; DG, dentate gyrus; U, unsusceptible; S, Susceptible.

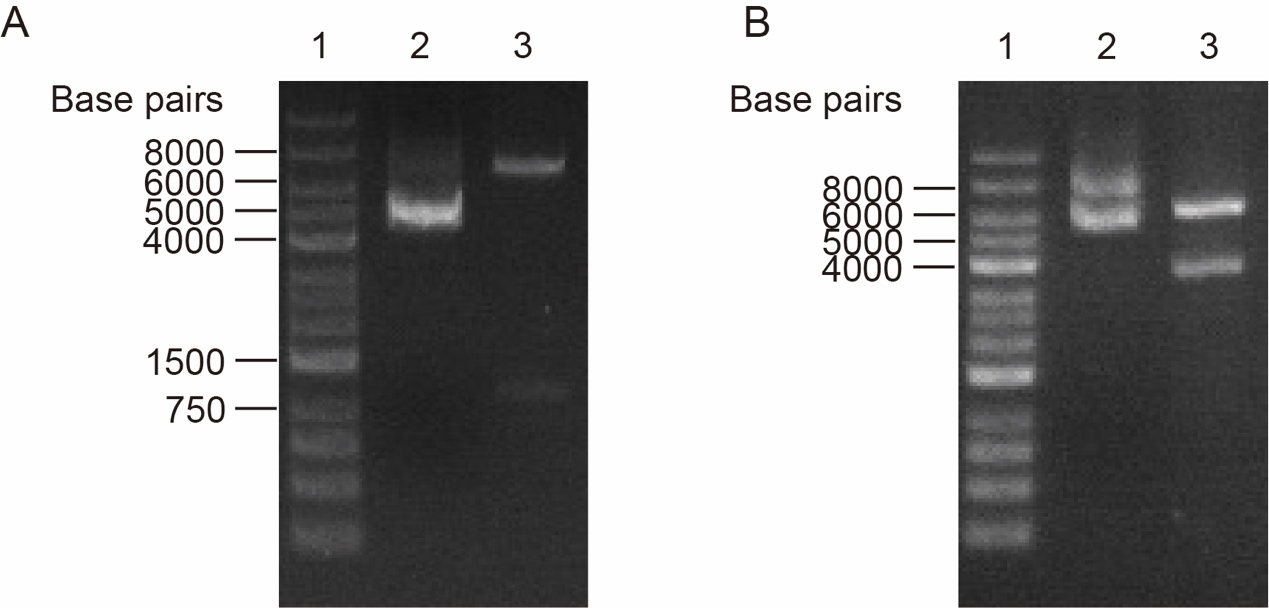

**Supplemental Figure 10. Verification for plasmid construction by PCR followed by restriction digestion. (A)** pBT3-STE-s1pr1 plasmid digested with XbaI-HindIII. **(B)** pPR3-C-itga2 plasmid digested with SalI-BamHI. Lane 1, 2, 3 in **(A-B)** represents DNA ladder, empty plasmid control and plasmid digested with restriction enzymes.

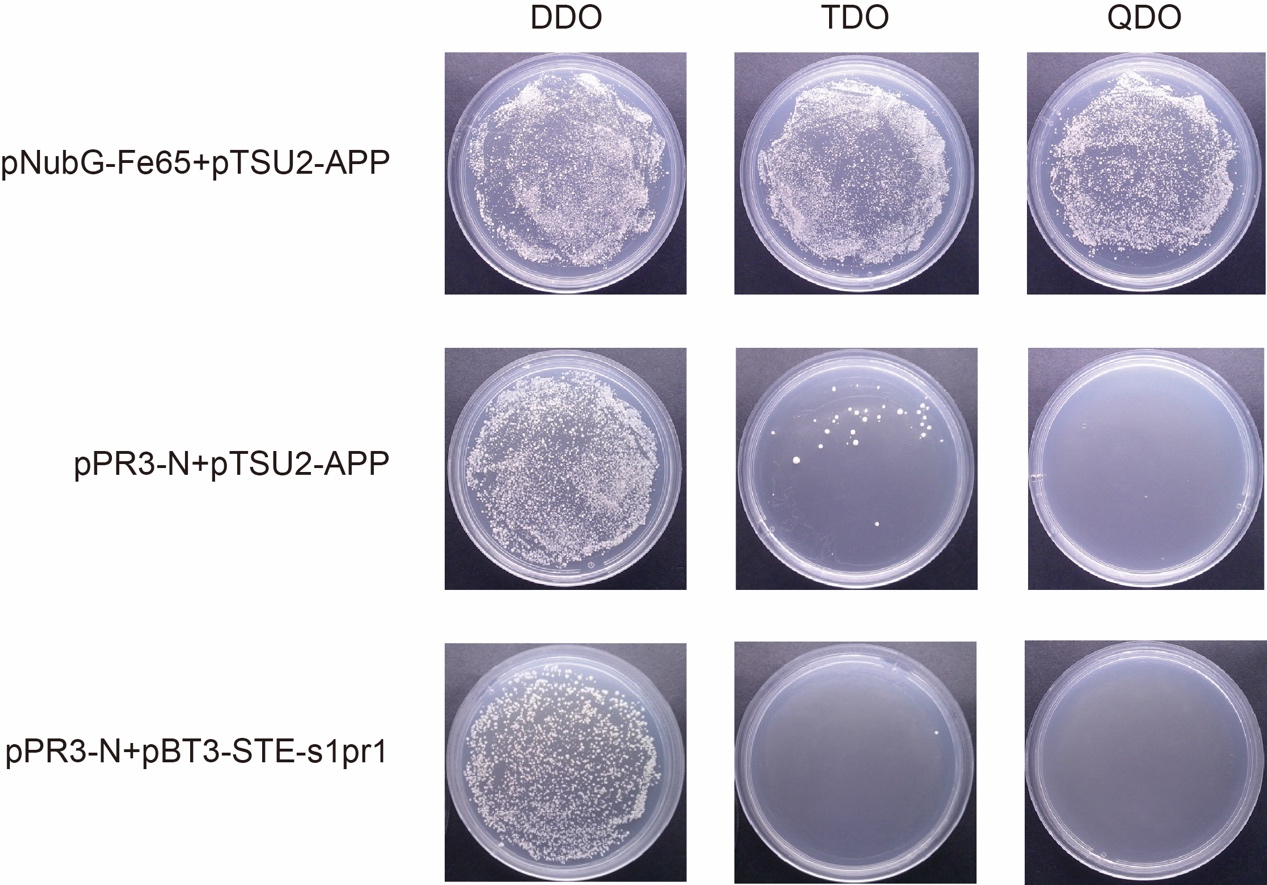

**Supplemental Figure 11. The auto-activation test.** Line 1, 2 and 3 represents a positive control (the pNubG-Fe65 and pTSU2-APP vector together), a negative control and the pPR3-N empty vector and the pBT3-STE vector with s1pr1 growing on the DDO (SD/-Trp/-Leu), TDO (SD/-Trp/-Leu/-His) and QDO (SD/-Trp/-Leu/-His/-Ade) plates, respectively.

**SUPPLEMENTAL TABLES**

**Supplemental Table 1. Virus vectors.**

| **Virus vector** | **Sequence** | **Vendor** |
| --- | --- | --- |
| rAAV- CaMKIIa-EGFP-shRNA(S1pr1)-WPREs | GCTCTACCACAAGCACTATAT | BrainVTA, Wuhan, China |
| rAAV-CaMKIIa-S1pr1-P2A EGFP-WPRE-hGH polyA | NCBI: NM_007901.5 | BrainVTA, Wuhan, China |
| rAAV-CaMKIIa-mCherry-shRNA(Itga2)-WPREs | GACCTCACAAACACCTTCAGA | BrainVTA, Wuhan, China |

**Supplemental Table 2. Chemicals.**

| **Chemicals** | **Vendor** | **Cat #** |
| --- | --- | --- |
| Triton X-100 | Sigma-Aldrich | 9036-19-5 |
| Normal Donkey Serum | Solarbio | SL050 |
| Meloxicam | Solarbio | M9840 |
| Dimethyl suifoxide | Solarbio | D8370 |
| SEW2871 | Aladdin | 256414-75-2 |

**Supplemental Table 3. Anti-bodies.**

| **Anti-bodies** | **Vendor** | **Cat #** | **Dilutability** |
| --- | --- | --- | --- |
| Anti-NeuN antibody [1B7] | Abcam | ab104224 | 1:500 |
| GFAP (GA5) Mouse mAb | Cell Signaling | 3670S | 1:500 |
| Anti-Iba1 antibody | Abcam | ab5076 | 1:500 |
| CaMKII alpha Monoclonal Antibody (6G9) | Invitrogen | MA1-048 | 1:500 |
| Anti-GAD67 Antibody, clone 1G10.2 | Sigma-Aldrich | MAB5406 | 1:500 |
| Donkey anti-Rabbit IgG (H+L) Highly Cross-Adsorbed Secondary Antibody, Alexa Fluor™ 488 | Thermo Fisher Scientific | **A-21206** | 1:500 |
| Donkey anti-Rabbit IgG (H+L) Highly Cross-Adsorbed Secondary Antibody, Alexa Fluor™ 594 | Thermo Fisher Scientific | A-21207 | 1:500 |
| Donkey anti-Mouse IgG (H+L) Highly Cross-Adsorbed Secondary Antibody, Alexa Fluor™ 488 | Thermo Fisher Scientific | A21202 | 1:500 |
| Donkey anti-Mouse IgG (H+L) Highly Cross-Adsorbed Secondary Antibody, Alexa Fluor™ 594 | Thermo Fisher Scientific | **A-21203** | 1:500 |
| Donkey anti-Goat IgG (H+L) Cross-Adsorbed Secondary Antibody, Alexa Fluor™ 488 | Thermo Fisher Scientific | A-11055 | 1:500 |
| Rac1 Polyclonal antibody | Proteintech | 24072-1-AP | 1:1000 |
| CDC42 Polyclonal antibody | Proteintech | 10155-1-AP | 1:1000 |
| ARP2 Polyclonal antibody | Proteintech | 10922-1-AP | 1:1000 |
| ARP3/ARP3B Polyclonal antibody | Proteintech | 13822-1-AP | 1:1000 |
| CD41/Integrin Alpha 2B Polyclonal antibody | Proteintech | 24552-1-AP | 1:1000 |
| GAPDH Monoclonal Antibody | Proteintech | 60004-1-Ig | 1:1000 |
| HRP-labeled Goat Anti-Rabbit IgG (H+L) | Beyotime | A0208 | 1:1000 |
